# Effect of ambient fluid rheology on oscillatory instabilities in filament-motor systems

**DOI:** 10.1101/2022.03.14.484323

**Authors:** Anupam Mishra, Joshua Tamayo, Arvind Gopinath

## Abstract

Filaments and filament bundles such as microtubules or actin interacting with molecular motors such as dynein or myosin constitute a common motif in biology. Synthetic mimics, examples being artificial muscles and reconstituted active networks, also feature active filaments. A common feature of these filament-motor systems is the emergence of stable oscillations as a collective dynamic response. Here, using a combination of classical linear stability analysis and non-linear numerical solutions, we study the dynamics of a minimal filament-motor system immersed in model viscoelastic fluids. We identify steady states, test the linear stability of these states, derive analytical stability boundaries, and investigate emergent oscillatory solutions and their properties. We show that the interplay between motor activity, aggregate elasticity and fluid viscoelasticity allows for stable oscillations or limit cycles to bifurcate from steady states. For highly viscous Newtonian media, frequencies at onset decay with viscosity *μ* as 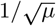. In viscoelastic fluids that have the same viscosity as the Newtonian fluid but additionally allow for temporary energy storage, emergent limit cycles are associated with higher frequencies. The magnitude of the increase in the frequency depends on motor mechanochemistry and the interplay between fluid relaxation time-scales and time-scales associated with motor binding and unbinding. Our results suggest that stability and dynamical response in filamentous active systems may be controlled by tailoring the rheology of the ambient environment.

## I. INTRODUCTION

Biology is replete with examples of rigid and semiflexible filaments, that singly or as aggregate bundles, interact with molecular motors in both intracellular and extracellular settings. For instance, actin-myosin systems are crucial for cell function, development and growth [1, 3–5], and for the transport of DNA and reproduction [2, 6–8]. Structured filament-motor systems seen inside eukaryotic cilia and flagella drive cellular motility [9], and are also crucial components of healthy respiratory and reproductive tracts [10–15]. In the synthetic realm, motor assays and reconstituted biofilament networks frequently feature molecular motors translating, deforming and bending filaments [16, 17, 34].

A characteristic feature of these filament-motor systems is the emergence of stable oscillations under favorable conditions. Examples are oscillatory instabilities in muscles and sarcomeres [18, 20, 21], oscillations in microtubule assays involving NCD motors [22, 23], the periodic undulations of eukaryotic cilia [7, 8], and spontaneous oscillations during asymmetric cell division that involve microtubules interacting with cortical force generators [24–26]. These oscillations may follow complicated periodic functions and are typically very sensitive and tunable. Often, the onset of these oscillations may be controlled by quenching the system by depleting the molecular motors of their energy source (ATP or Ca^2+^). Another crucial ingredient in these systems is that they are not just open systems with continuous energy input, but are also usually highly non-linear. All these suggest that the onset of these oscillations may be interpreted as instabilities to a base state triggered by system changes.

Biofilaments and motors usually inhabit fluidic or a gel-like media with highly non-Newtonian properties. The complex fluid environment not only provides a way for the transfer fuel but also directly interacts with aggregates by exerting time-dependent stresses. For instance, a filament moving in a Newtonian fluid feels viscous drag stresses exerted by the ambient medium. A filament moving in viscoelastic environments will be subject to more complicated time dependent forces with both dissipative (viscous) and non-dissipative (elastic) components. Recent work on extracellular cilia that enables whole-cell locomotion [9, 28] suggests a strong influence of the viscosity and complex rheology of the ambient fluidic medium. Sketched in Figure 1(a) is a schematic of experiments from [9] that depict variation in ciliary beat frequency in a model Newtonian (red) and polymeric (blue) fluid. The oscillations in the viscoelastic polymeric fluid are significantly higher than in a Newtonian fluid with the same viscosity. In the human mucociliary tract in the presence of disease or ciliary dysfunctions such as cystic fibrosis (CF), primary ciliary dyskinesia (PCD) or chronic obstructive pulmonary disease (COPD), the mucus layer submerges cilia in an extremely viscoelastic environment that impacts their beating [10, 12–14] Thus the effect of the rheology of the ambient fluid needs to be understood in order to predict the dynamical response in these biological filament-motor systems. There is however a significant gap in our understanding of these aspects since experimentally it is difficult to independently vary fluid viscosity and elasticity.

**FIG. 1.**
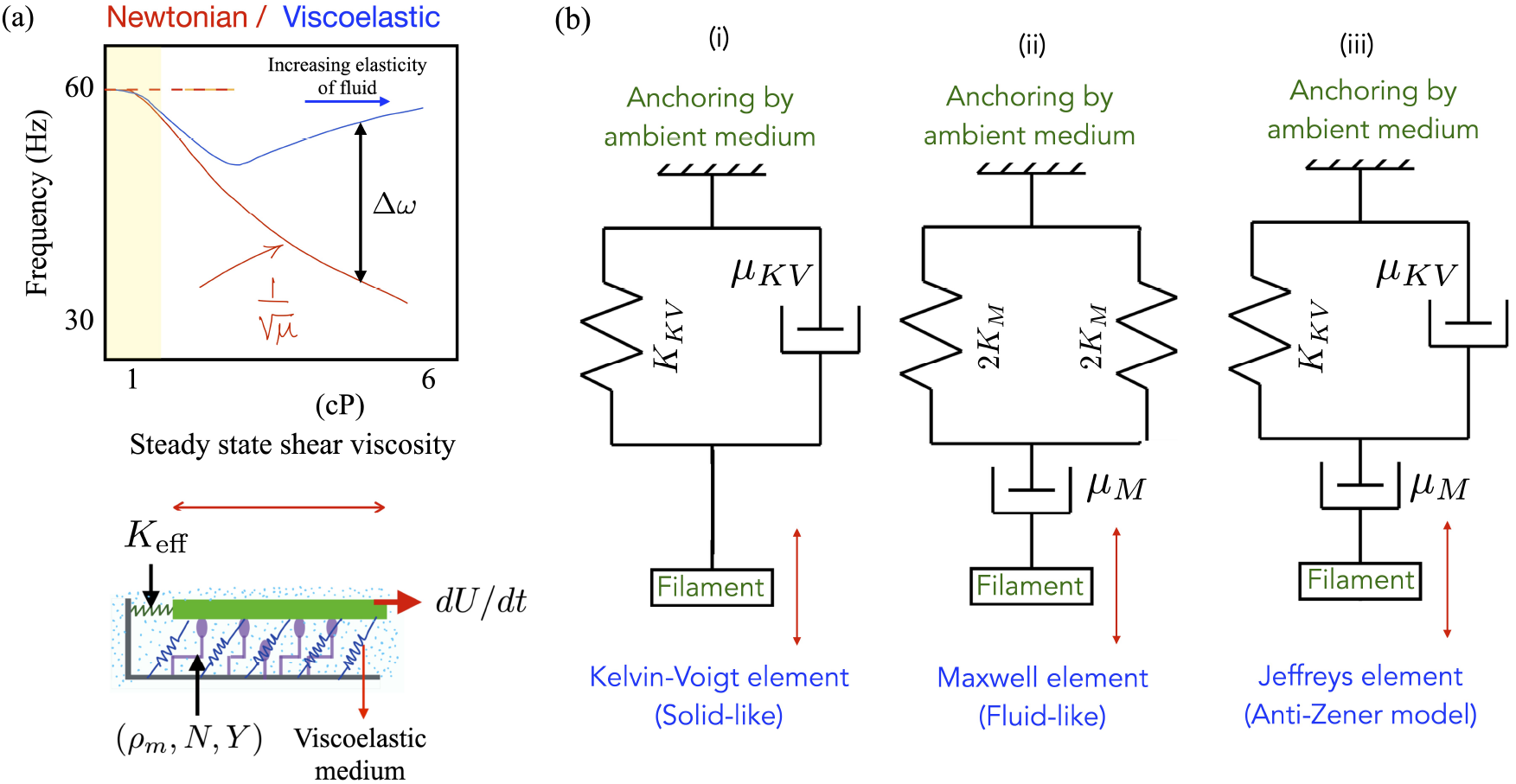
(a) (Top) A sketch of experimental observations of ciliary frequencies on bi-ciliated alga *Chlamydomonas reinhardtii* moving in model viscoelastic fluids [9] made by adding polymer to a Newtonian solvent. The red curve demonstrates the functionality seen in a purely Newtonian fluid. The blue curve shows how the frequency in polymeric fluids is much higher than in Newtonian fluids with the same viscosity. (Bottom) Schematic of the animated motor-filament segment that forms the basis of the minimal motor-filament system. Active motors (purple) may be in one of two states - attached to the segment (in green) or detached. Passive cross-linkers are shown in blue and are treated as linearly elastic springs. The resistance to motion of the test segment exerted by its neighbors are combined into an effective elastic resistance (spring constant *K*_eff_). The ambient medium in which the assembly is embedded is viscoelastic. In the minimal model this medium may be treated as a Maxwell medium or as a Kelvin-Voigt medium. (b) Representations of the viscoelastic ambient media considered in this article. (a) A Kelvin-Voigt material, (b) a Maxwell fluid, and (c) a generalized Jeffrey element.

Abstracting the ingredients crucial to the examples of actively driven motor-filament systems allows us to identify three main ingredients that may control system-level mesoscale and macroscale dynamics - a) passive elasticity, from intrinsic filament-motor elasticity, b) rheological properties of the embedding medium, and c) associated motor mechanochemistry and kinetics. Here, we will combine these elements to build a simple phenomenological model for a filament-motor aggregate interacting with a viscoelastic medium. To enable analytical progress, we assume the ambient fluid to respond in one of three canonical ways – as a Newtonian fluid, as model Kelvin-Voigt gels or as a model Maxwell medium. Using this minimal model we investigate the onset, stability and control of oscillations and address associated questions: 1) how do variations in the fluid induced mechanical stresses affect the manner in which the internal biomechanical state of the molecular motors in the aggregate is modulated, 2) how do these modulations impact the onset of oscillations and when may they stabilize an otherwise unstable system, and 3) how does fluid rheology affect the frequency of emergent oscillations and their amplitude?

Biological filament-motor systems are much more complex than the minimal model analyzed here, and the emergence of oscillations in a complex biological system almost always depends crucially on the dynamic properties of the interacting components *and* their collective behaviors. Nonetheless, minimal models offer an essential tool for understanding basic biophysical mechanisms. In this sense, the simple minimal model analyzed here using model fluids represents an important step towards more complete computational models.

## II. MINIMAL MODEL

Our model system is shown in Figure 1(a) (bottom pane). We model the filament or filament bundle – referred to henceforth as just (composite) filament – as a rigid sheet. In the reduced setting analyzed here, we assume that the sheet has length *ℓ*_*A*_, thickness *b*_*A*_ ≪ *ℓ*_*A*_ and lateral extent *w*_*A*_ satisfying *ℓ*_*A*_ ≫ *w*_*A*_ ≫ *b*_*A*_. The view shown in the figure is from the side, the axial dimension is the length and the lateral dimension into the paper (corresponding to the lateral extent, *b*) is treated as a neutral direction. The area of the segment that faces the underlying substrate is *ℓ*_*A*_*w*_*A*_. Of course, biofilaments or biofilament aggregates may have slender cylindrical geometries; the area *ℓ*_*A*_*w*_*A*_ is thus to be interpreted as the projected area of the aggregate that interacts with molecular motors.

The filament is held a fixed distance away from an underlying flat substrate by passive, permanent linear springs (blue springs) with areal density *ρ*_p_, stiffness *K*_p_, and equilibrium length *ℓ*_p_. These passive springs resist shearing deformations and lateral translation of the filament. Additionally, the filament interacts with a collection of active motor proteins shown in purple. These motors are grafted to the lower plate with an areal density *ρ*_m_. We treat these motors as active linear elastic springs with spring constant *k*_m_ with an equilibrium rest length *ℓ*_m_. The tail of each motor is attached to the substrate; their heads meanwhile can periodically adhere to the filament and generate forces displacing the filament laterally plate to translate laterally as shown. For ease of analysis without loss of generality, we set *ℓ*_m_ = *ℓ*_p_ = 0. We choose the inter-motor spacing to be small enough, and densities high enough that a continuum description of filament-motor interactions suffices.

Our focus here is on generic features and physical mechanisms of oscillations in molecular motor assemblies embedded in complex fluids. To aid in the analysis, we model motor-filament interactions using two-state crossbridge models [8, 18, 21]. Each motor is is either attached state or detached. Detached motors can attach to the filament in a forward-leaning position with specified attachment probabilities. They then quickly undergo a conformational change, which makes them strained. As the filament moves, so does the motor and thus results in the extension of the motor and thus extension of the internal spring (with spring constant *k*_*m*_). Attached motors can detach any time, and the statistics of this process is embodied via microscopic strain dependent detachment probabilities. The transitions between attached and detached states – the kinetics of the mechanochemical cycle – are thus determined by motor kinetics, and specifically the attachment and detachment probabilities (rates).

Detailed microscale equations for motors such as these have been derived and rationalized earlier by others and by us [18, 35]. Briefly summarized, motor kinetics may be described by a set of population balances relating the attached and the detached probability densities to the attachment and detachment fluxes via microscopic transition rates. For simplicity we neglect motor diffusion, and consider the noiseless mean-field limit, where the number of motors (and the density) is large (high) enough that fluctuations in the time average of the density of attached motors are small compared to the mean value. When the distribution in extension of attached motors is sharply peaked about the typical (average) length, transients to this distribution occur over times very small compared to the averaged macroscopic time scale. The extension of attached motors is then peaked about the mean value with small deviations from the mean. We assume that detached motors relax to a delta function with the change occurring instantaneously. Coarse-graining the microscale dynamical equations provides a mean-field continuum description for the (mean) attached motor fraction, and the mean attached motor extension. Motor kinetics is captured by the mean-field attachment rate *ω*_on_, and the mean-field detachment rate *ω*_off_.

## III. DIMENSIONLESS COUPLED EQUATIONS

Following the physical picture described in §2, we derive equations governing the dynamics for an elastically constrained filament-motor assembly embedded in a viscoelastic fluid (detailed derivation in Appendix Section A). To enable analytical progress, we assume that the rheology of the embedding fluidic medium behaves (a) a Newtonian fluid, (b) a Kelvin-Voigt material, or as a (c) Maxwell medium (Supplementary Information §1). Based on these, the filament interaction with the ambient fluid may be calculated analytically. This process results in equations (1)-(4) constitute the (scaled) nonlinear ODE’s governing the evolution of the motor-filament system embedded in the viscoelastic medium.

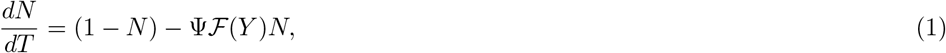

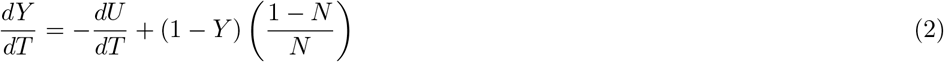

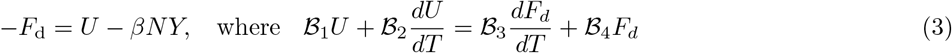

with dimensionless parameter *β* quantifying motor density while dimensionless parameters ℬ_1_ to ℬ_4_ quantifying the viscoelasticity of the fluid and fluid specific relaxation times.

Following previous work, we model motor kinetics as responding to the deformation experienced by attached motors [1, 2, 18, 19]. Attached motors behaving as slip bonds with a strain dependent detachment probability. This functionality is encoded in the detachment rate 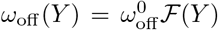 that modifies the bare detachment rate by a multiplicative factor (ℱ*Y*). For the analytical investigations, we restrict (ℱ*Y*) to functions that is at least twice differentiable but otherwise general. For plotting, we will use the following forms

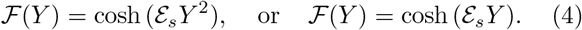

Equations (1)-(4) admit a stationary state given by the set (*N, Y, U*) = (*N*_0_, *Y*_0_, *U*_0_) where these satisfy

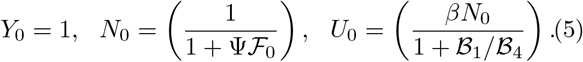

In equation (5), the dimensionless parameter *ε*_*s*_ quantifies the characteristic force attached motors can with-stand before detachment. Note that the ratio ℬ_1_/ℬ_2_ compares the motor time scale with the relaxation time scale of the Kelvin-Voigt medium. Dimensionless parameter ℬ_3_ is similarly a ratio of motor time-scales to fluid relaxation times scales for the Maxwell fluid. Physical interpretations of variables (*U, N, Y*) and that of dimensionless parameters are summarized in Tables 1 & 2.

**TABLE I.** Motor-related parameters and scaled variables in the minimal model.

| Symbol | Definition | Interpretation |
| --- | --- | --- |
| $\mathcal{A}$ | $d_m$ | Typical length scale of motor displacement and strain |
| $\mathcal{E}_s$ | $k_m d_m / k_B T$ | Strain dependent force scale for motor detachment |
| $\beta$ | $\rho_m k_m / (\rho_p k_p + \bar{K}_{\text{eff}})$ | Scaled motor density and a measure of active elasticity |
| $\Psi$ | $\omega_{\text{off}}^o / \omega_{\text{on}}^o$ | Ratio of motor kinetic constants and related to motor duty ratio |
| $N(t)$ | $\rho_a / \rho_m$ | Ensemble averaged fraction of total motors attached |
| $Y(t)$ | $Y^* / \mathcal{A}$ | Ensemble averaged (mean) motor extension |
| $U(t)$ | $U^* / \mathcal{A}$ | Scaled filament extension (strain) |

**TABLE II.**
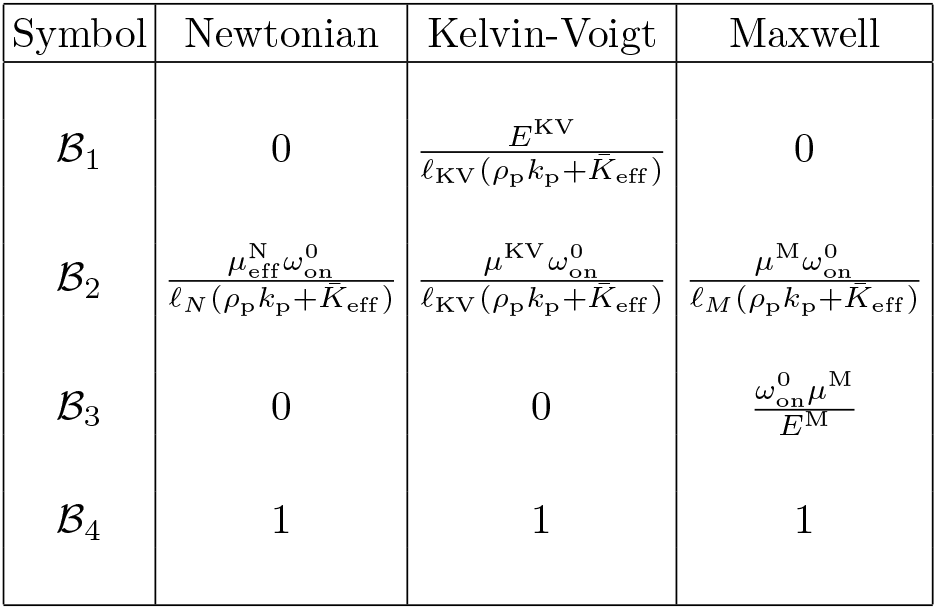
Dimensionless parameters quantifying the ambient medium stress on moving filament. Here, *ℓ*_*A*_ and *w*_*A*_ are the length and width of the filament aggregate interacting with motors and allow one to go from the stress to a force description. Displacements are scaled with 𝒜. We assume a continuum description is valid and that strains in the fluid/medium network may be appropriately defined using elastic and viscous moduli. We consider three cases: (a) a Newtonian fluid, (b) Kelvin-Voigt liquid-like viscoelastic medium, and (c) Maxwell solid-like medium. Models (b) and (c) serve as limiting cases of the more general linear Jeffreys’ model that belongs to the class of anti-Zener models in classical viscoelasticity. Note that the ratio ℬ_2_/ℬ_1_ for the Kelvin-Voigt model is a ratio of rheological to motor kinetics time-scales, just as parameter ℬ_3_.

In our model, bending is ignored and the filament composite/aggregate is considered rigid and inextensible. While our model is not directly applicable to ciliary oscillations, using some numerical estimates relevant to filaments and motors in cilia provides understanding of the magnitudes of the parameters in the model. Our model also may allow us to interpret how oscillations may arise within and along an initially straight passive cilium. Localized elastically weak regions may result in cilia fragments driven by arrays of dynein moving against their neighbors. In this scenario, we estimate *k*_*m*_ ∼ 10^−3^ N/m, the effective linear density *ρ*_*m*_*w*_*A*_ ∼ *O*(10^8^) m^−1^, *w*_*A*_ ∼ 40 − 60 nm, *k*_N_ ∼ 16 − 100 pN *μ*m^−1^ and *w*_*A*_*ρ*_N_ ∼ 10^5^ − 10^7^ m^−1^ [1, 27]. The value of *K*_eff_ depends on the material. Microtubules are relatively stiff with large persistent lengths and the Young’s modulus *E* ∼ 1.2 GPa providing the stiffness per length *K*_eff_ ∼ *Gw*_*A*_. For microtubules driven by dynein, previous studies report a persistence length ∼ 5mm at room temperature. For actin driven by myosin, ∼ 16 *μ*m at room temperature. These length scales provide an upper bound for *ℓ*_*A*_.

## IV. RESULTS AND DISCUSSION

### A. Dynamics of the unconstrained system

Before analyzing the dynamics of the constrained filament-motor system, we briefly address the dynamics of an unconstrained system. Accordingly, we set *ρ*_*p*_ = 0, *K*_eff_ = 0 and study the dynamics of the animated filaments on a rigid surface on which motors are grafted, as in motility assays. Short fragments in these assays are typically rigid enough that they move without bending. Long fragments on the other hand are susceptible to buckling instabilities as previous studied experimentally and analytically [29–34, 36] in these and similar systems involving driven active filaments.

Here, we invert the problem and study the dynamics of the motor variables (*Y, N*) as functions of the speed *Z* = *dU/dT*. In other words, we restrict ourselves to time scales where the dynamics of the motor kinetics is slaved to the speed of the filament. The speed *Z* may be envisioned as an independently controlled variable; we then enquire how steady values of *N* and *Y* depend on *Z* and if these solutions are linearly stable. Equation (10) is irrelevant in this limit, and equations (1) and (2) become

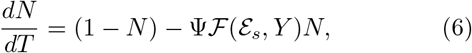

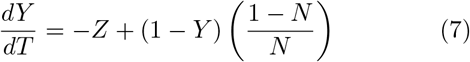

When *Z* = 0, equations (6) and (7) admit steady state solutions (*N*_*s*_, *Y*_*s*_) given by

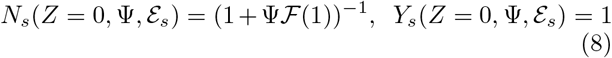

For each value of *Z* ≠ 0, other steady solutions different from (8) exist. To study both the steady solutions and their stability, we employ an in-house numerical continuation implemented in Matlab^@^. Treating *Z* as the free variable, we plot steady solutions to equations (6) and (7) and investigate the stability of the solutions for various values of Ψ and _*s*_ and for different functional forms of ℱ the detachment function. Figure 2 shows an example case where we keep *ε*_*s*_ and Ψ the same while using two forms of ℱ. Solid curves (maroon and blue) represent stable solutions while dashed curves are unstable solution branches. Turning points, where these branches meet, are shown as circles.

**FIG. 2.**
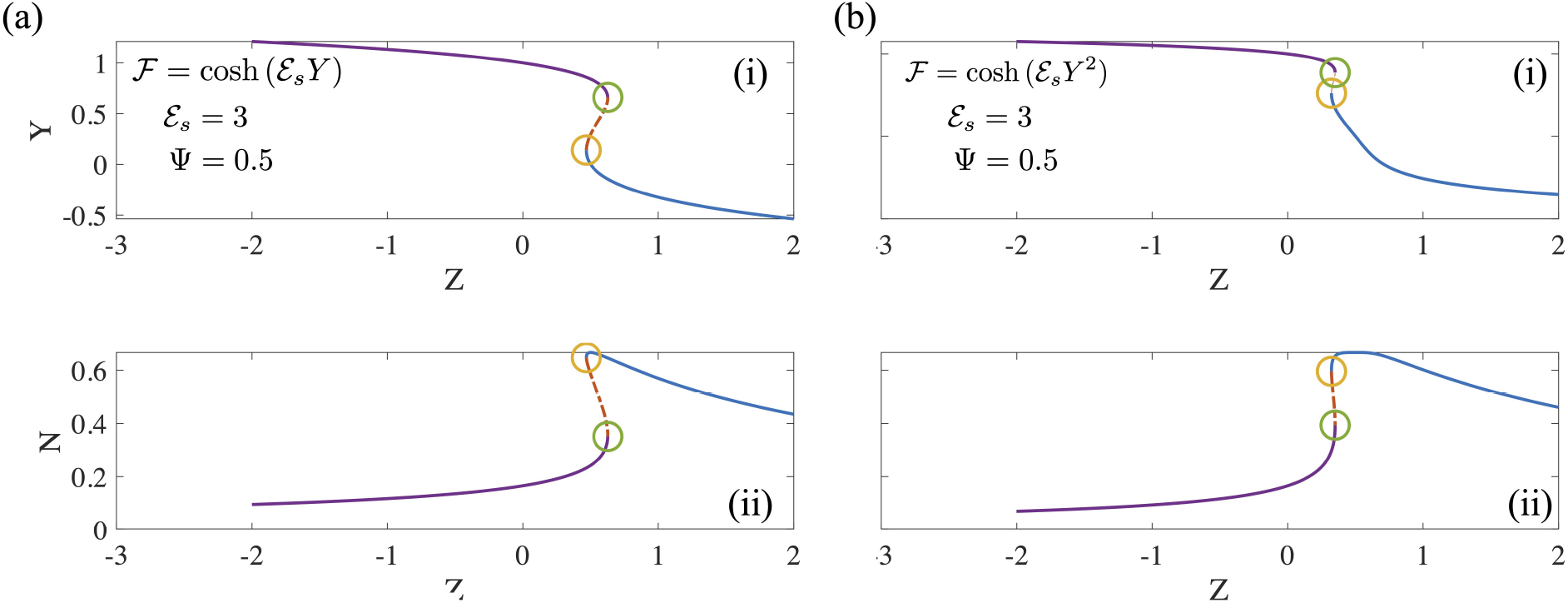
The mean motor extension *Y* and fraction of motors that are attached *N* as a function of *Z* (steady solutions to equation (6,7)) for two forms of the detachment function (a) ℱ = cosh (*E*_*s*_*Y*) and (b) ℱ = cosh (*ε*_*s*_*Y* ^2^).

Numerical investigation reveals that the existence and extent of the unstable region depends on the form of the detachment curve. When the detachment function is constant or a decreasing function of extension *Y*, no unstable regions exist. In these cases, typically each value of *Z* is associated with a unique value of *N* and *Y*. However when the detachment function is an increasing function of *Y*, some parts of the steady state solutions becomes unstable as seen in Figure 2. This is manifest clearly in bistability, evident for instance in the curves of *N* vs *Z* with three possible values of Z (two stable, one unstable) corresponding to a single value of *Z*. A value of *N* = 0.4 in Figures 2(a)-(ii) and 2(b)-(ii) for instance is unstable and is susceptible to disturbances that can push the state to either the upper branch or the lower branch each with the same value of *Z*. The origin of this instability is the emergence of effective negative spring constants due to motor activity; such features have been studied previously [18].

### B. Analytical results: Oscillatory instabilities and emergent frequencies

The question that arises naturally next is how preventing constant speed solutions by allowing *K*_eff_ *>* 0 impacts this response. To answer this question, we turn next to the stationary base state of the full equations (1-4) and the linear stability of the base state given by (5). Mathematical details of the linear stability analysis are provided in Appendix B. We find that the base state exhibits linear instability for certain regions in parameter space. We derive analytically expressions for the locus of these critical points – the neutral stability curves (presented in Supplementary Information §2). Classical linear stability analysis shows at these critical points, stable oscillatory solutions emerge via the Hopf-Andronov-Poincare bifurcation [37]. Limit cycles emanate at these points with fixed frequencies and zero amplitude. We calculate the frequencies of these emergent solutions analytically and investigate their dependence on dimensionless parameters defined earlier. Predictions of our linear stability analysis are confirmed by full non-linear solutions of the equations.

#### 1. No drag from medium

Consider the simplest degenerate case where the medium does not exert any forces on the motor-filament system. In this case, we have ℬ_1_ = ℬ_2_ = ℬ_3_ = 0 and ℬ_4_ = 1. The equation for the growth rates is a quadratic equation for this degenerate case of the form *bσ*^2^ + *cσ*+ *d* = 0 where *σ* is the linear growth rate of infinitesimally small disturbances. For oscillatory instabilities, we require that *c* < 0 (so the growth rate, *σ* is non-negative); with this holding, *c* = 0 defines the locus of critical points. The neutral stability curve follows

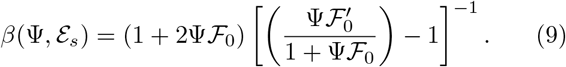

For constant Ψ and *ε*_*s*_, equation (9) provides the critical value of *β* above which one can expect oscillatory limit cycles to bifurcate from the stationary steady state. Note that an essential condition here is 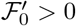 and that the magnitude be large enough for the equation to be satisfied. Formally, we require ℱ to be a monotonically increasing function of *Y* consistent with previous work (see [2] for instance) in similar systems. The periodic solutions at onset have frequency

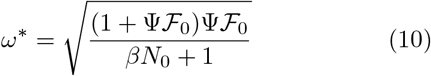

#### 2. Newtonian fluid

For the simplest drag-producing medium - a Newtonian fluid - we have ℬ_1_ =ℬ _3_ = 0, ℬ_2_ > 0 and ℬ_4_ = 1 with specific forms listed in Table 1. The force balance reduces to

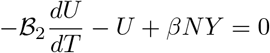

with first two left hand side terms being resisting (passive) forces and the the third term the driving (active) force; the right hand is zero here due to the absence of inertia as explained earlier. The frequency of the bifurcating limit cycles at onset is given (Appendix B) by

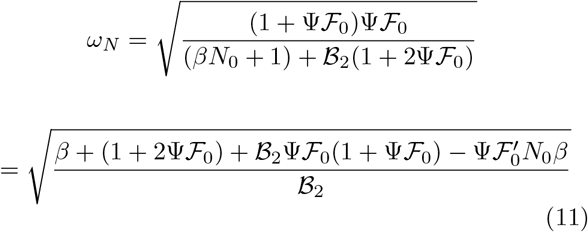

Combining (10) and (11) yields

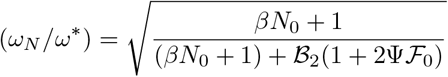

and so,

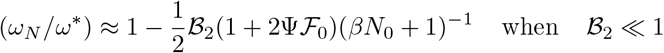

while

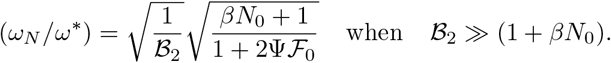

Thus for very high viscosity *μ*_eff_, the frequency scales as 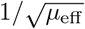. From the analytical expressions for the neutral stability curve we deduce that for oscillatory instabilities to exist we require that 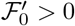 and be a suitably large number. For large ℬ_2_ such that ℬ_2_ ≫ (*βN*_0_ + 1), the critical value of *β* for onset of instability *is larger* than for the case of no drag and is consistent with expectation, since extra viscous drag implies higher dissipation rates. At constant motor properties – that is, at constant values of Ψ and *ε*_*s*_ and with the form of ℱ (*Y*) held fixed – the only way to enable this is by having a larger number (density) of active motors.

#### 3. Kelvin-Voigt medium

Here, the drag force depends on the displacement and the rate of displacement. Thus we have ℬ_4_ = 1, ℬ_1_ > 0, ℬ_2_ > 0 while ℬ_3_ = 0. In this case again, oscillatory instabilities are possible at critical points provided 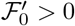. The frequency of the bifurcating limit cycles at onset is obtained as

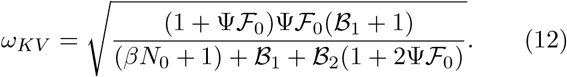

For fixed motor parameters and fixed value of ℬ_2_, the frequency in a KV medium is thus higher than the purely viscous medium (with the same viscosity/dissipative components, ℬ_2_) consistent with experiment. The increment comes from the elastic component of rheology of the fluid but is augmented by the motor activity parameter Ψ and the strength of the detachment function but *not by its slope*. To compare with the two limiting cases analyzed earlier, we note that

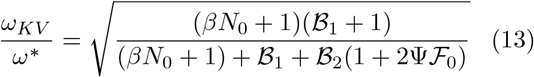

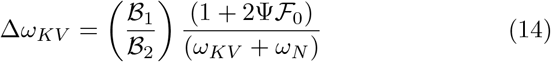

where we have defined the shift in frequency relative to a purely Newtonian fluid, Δ*ω*_*KV*_ ≡ (*ω*_*KV*_ − *ω*_*N*_).

#### 4. Maxwell medium

For the Maxwell medium, the displacement rate is linearly coupled to the drag force and the rate of change of the drag force. We have for this case, ℬ_1_ = 0, ℬ_2_ > 0, ℬ_3_ > 0 and ℬ_4_ = 1. From an estimation of constants *a* − *d* (Supplementary Information § 2), we deduce that the cubic equation for the growth rate cannot be zero (ie. real and imaginary parts both zero). This rules out the possibility of a fold-Hopf bifurcation, that is a fold bifurcation coincident with the Andronov-Hopf-Poincare bifurcation.

However we can still satisfy conditions for the existence of an oscillatory instability. We therefore look for a subset of critical points for which exactly 1 real negative root (*σ*_1_) and a complex conjugate pair (±*iω*) with zero real part (*σ* = 0) exists for the cubic. The neutral stability curve is defined by the locus of points that simultaneously satisfy *ad* − *bc* = 0. The frequency at onset is given by

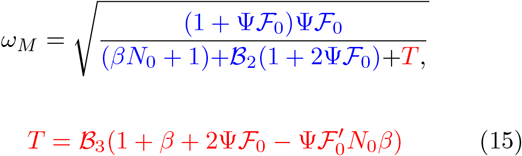

The terms in blue on the right hand side of (15) are terms that appear in the equation for the emergent frequency for a Newtonian fluid while the term in red is the extra term that appears with the Maxwell medium. Since ℬ_3_ > 0, the overall sign of the term depends on the magnitude of the term 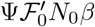 that has a negative sign ahead of it. We have deduced earlier that for oscillations to exist we require that the detachment function be an increasing function of *Y* evaluated at the steady point *Y*_0_. Combining these observations, we deduce that the frequency in a Maxwell medium can be higher than the Newtonian medium provided the term in red is negative. In fact the magnitude of this term (when negative in sign) can be readily modified two ways. First, for fixed ℬ_3_, the magnitude can be adjusted by solely changing the form of the detachment function (through 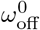 and/or *ε*_*s*_ and *independent* of the rheology parameters and the motor attachment rate. Second, for fixed motor mechanochemistry as long as the term in the parenthesis is negative, increasing ℬ_3_ will increase the frequency of oscillations.

Before concluding this section, we address the biophysical origin of the oscillations. The simplest scenarios – the case of no fluid, and that of a Newtonian fluid – provide hints as to the destabilizing mechanism. Assume that the base state identified in (5) is stable. Linearizing (5) about the base state provides (see also Appendix B, equations (28)-(29)) in frequency space a relationship between the displacement *U* and an effective compliance that includes contributions from the active motor-based interactions with the filament as well as the fluid response. When there is no drag/resistive force at all from the medium, we find that the instability arises when the active compliance has a negative effective elastic coefficient i.e, active motor induced elasticity overcomes the passive elastic resistances. This provides a mechanism by which *system-level* oscillations are initiated. Meanwhile the dissipative components of the system - i.e, viscous effects - limit the amplitude of the oscillations. Thus stable limit cycles can emerge. Interestingly no fluid or external medium is needed to initiate and sustain oscillations in this model system.

We have three independent parameters here that determine if oscillations exists - *β*, Ψ and *ε*_*s*_. In general, we need the motors to detach so that the filament can slide back and forth. Very high motor attachment rates with no detachment also impedes oscillations by increasing the effective shear stiffness of the filament. Thus there is a lower limit for oscillations in terms of Ψ. For very high Ψ, the linear stable phase-space occupies larger extent. Similarly, there is a minimum value of *ε*_*s*_ below which oscillations cannot be sustained – this is because motor detachment is not sufficient to sustain reversal of motion of the filament. Very high values of *ε*_*s*_ on the other hand imply that motors detach almost as soon as they attach. Finally, there is also a minimum value of *β*, or equivalently a minimum density of active motors, needed to initiate oscillations.

### C. Comparison to non-linear solutions and discussion

The linear stability analysis in Section 4.1-4.2 provides information on the stability boundaries for the stationary steady state in (5) and also on the frequencies of limit cycles at the critical points on these boundaries. The actual frequency when one is far away from the neutral stability curve will of course differ from the values at criticality. Nonetheless, the emergent frequencies provide a good estimate and qualitative understanding of how the different biophysical parameters impact the frequency. Specifically, we are able to analytically calculate how motor density and elasticity (*β*), motor kinetics (Ψ and *ε*_*s*_), and fluid rheology (parameters ℬ_1_ to ℬ_3_) impact the neutral stability curve and frequencies. The predicted trends are line with experimental observations of more complicated systems with both sliding and bending deformations such as that shown in Figure 1(a). Making a direct comparison isn’t possible, but it is heartening to note that significantly higher frequencies (increase of up to a factor of 10) can be achieved by varying the rheological parameters of the model fluids, such as by keeping viscosity via ℬ_2_ fixed while changing ℬ_1_ in the Kelvin-Voigt model and ℬ_3_ in the Maxwell model.

Our analysis also suggests that smooth modulation of the frequencies may be implemented once past the critical point but linear stability alone is insufficient to validate this. Furthermore the linear stability analysis does not show how the amplitude of the oscillations changes with parameters. The amplitude is dominated by nonlinear terms. Thus to understand how non-linearity affects frequencies and amplitudes far from critical points, we solved equations (1)-(4) numerically using Matlab^@^. Equations (1)-(4) constitute a stiff set of equations especially in the limit of small ℬ_2_ or large ℬ_3_ and hence the suite for stiff equations was employed. Sample time-dependent solutions are shown as follows: (a) no drag case - Figure 3(b), for (b) Newtonian fluid sample solutions are shown in Figures 4(a) & 4(b), for the Kelvin-Voigt material Figures 5(e) & 5(f) demonstrate important features of the full limit cycles, and finally more elaborate results for the Maxwell fluid are shown in Figure 6.

**FIG. 3.**
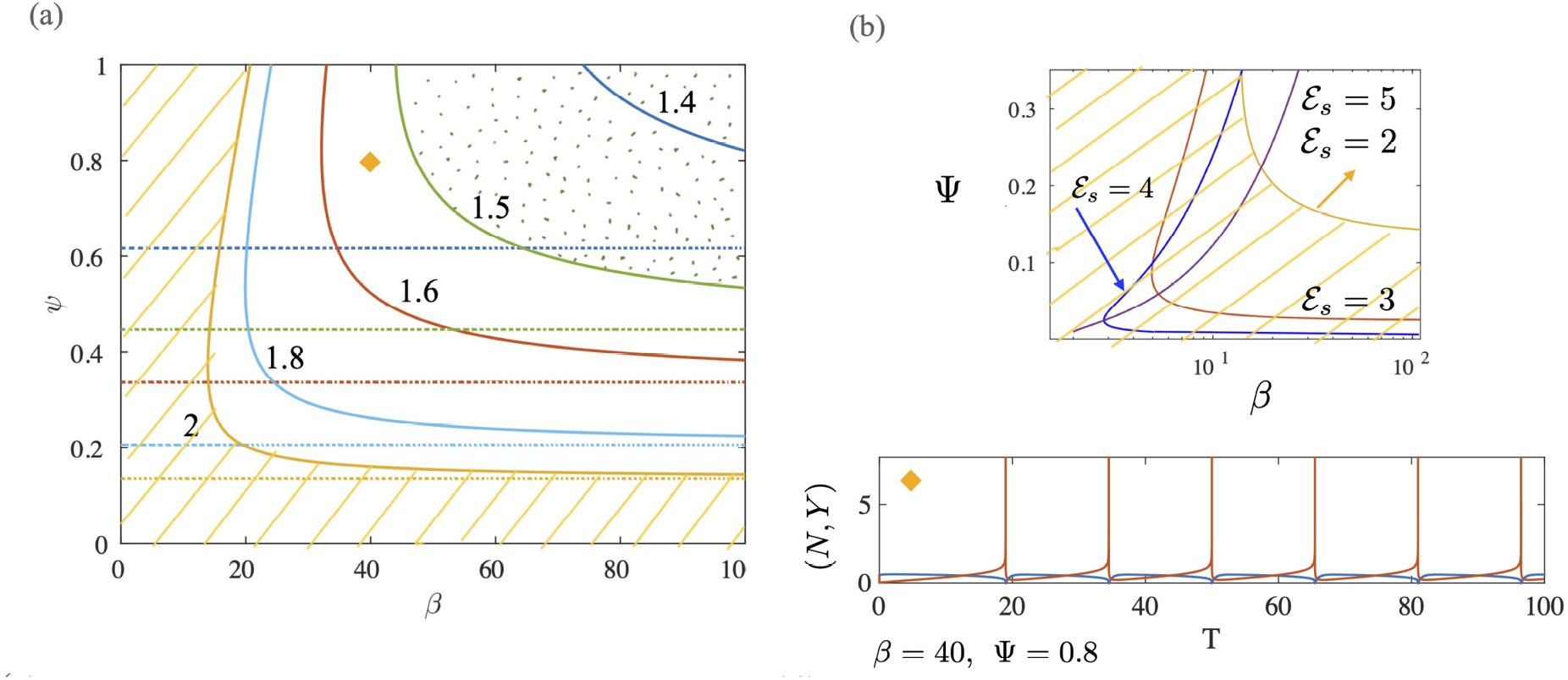
(a) Neutral stability curves separating linearly stable from unstable regions are shown for the no drag case. Plotted as solid curves are results for different values of *ε*_*s*_ (as indicated, from 1.4-2). The dash-dot lines are the limiting values of Ψ below which the base state is always stable. The shaded yellow region indicates the stable region for Ψ = 2. The dotted green region indicates linearly unstable space susceptible to oscillatory instabilities for Ψ = 1.5. The yellow rhombus point indicated in (a) is unstable for some parameter values, as demonstrated by full non-linear solutions when Ψ = 2 (shown alongside). (b) For even larger values of *ε*_*s*_ > 2, the curves shift rightward as Ψ increases. In all these cases, ℱ (*ε*_*s*_, *Y*) = cosh (*ε*_*s*_*Y*).

**FIG. 4.**
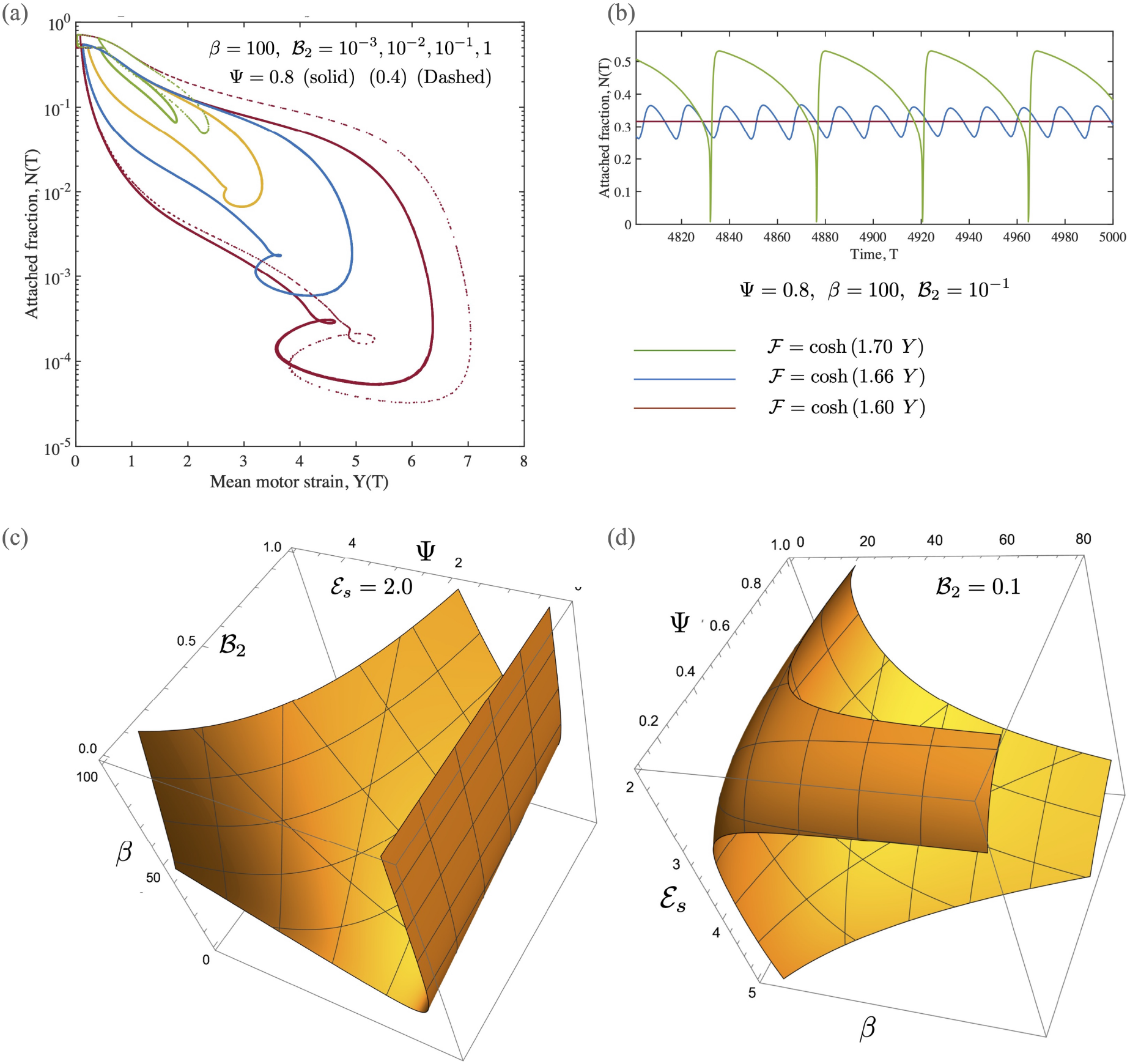
(a) Phase-plots over complete cycles obtained from fully non-linear solutions for the Newtonian case are shown. The detachment function is cosh (2*Y*) and the motor activity parameter (density) *β* = 100. We note that as ℬ_2_ increases from 10^−3^ to 10^−1^, the amplitude of oscillations decreases in sync with smaller variations in *N* and in *Y* over a cycle. (b) When motor density and kinetic parameters (*β* and Ψ) are kept constant, the value of *ε*_*s*_ still controls if oscillations can ensure. Shown here are non-linear solutions – at ℬ_2_ = 10^1^ – demonstrating the quenching of oscillations as *ε*_*s*_ is reduced from 1.7 to 1.6. (c,d) Neutral stability curves shown here for the Newtonian case when ℱ = cosh (*ε*_*s*_*Y*). Typically there are critical values of Ψ (for fixed *β*), or of *β* (for fixed Ψ) below which the base state is always stable.

**FIG. 5.**
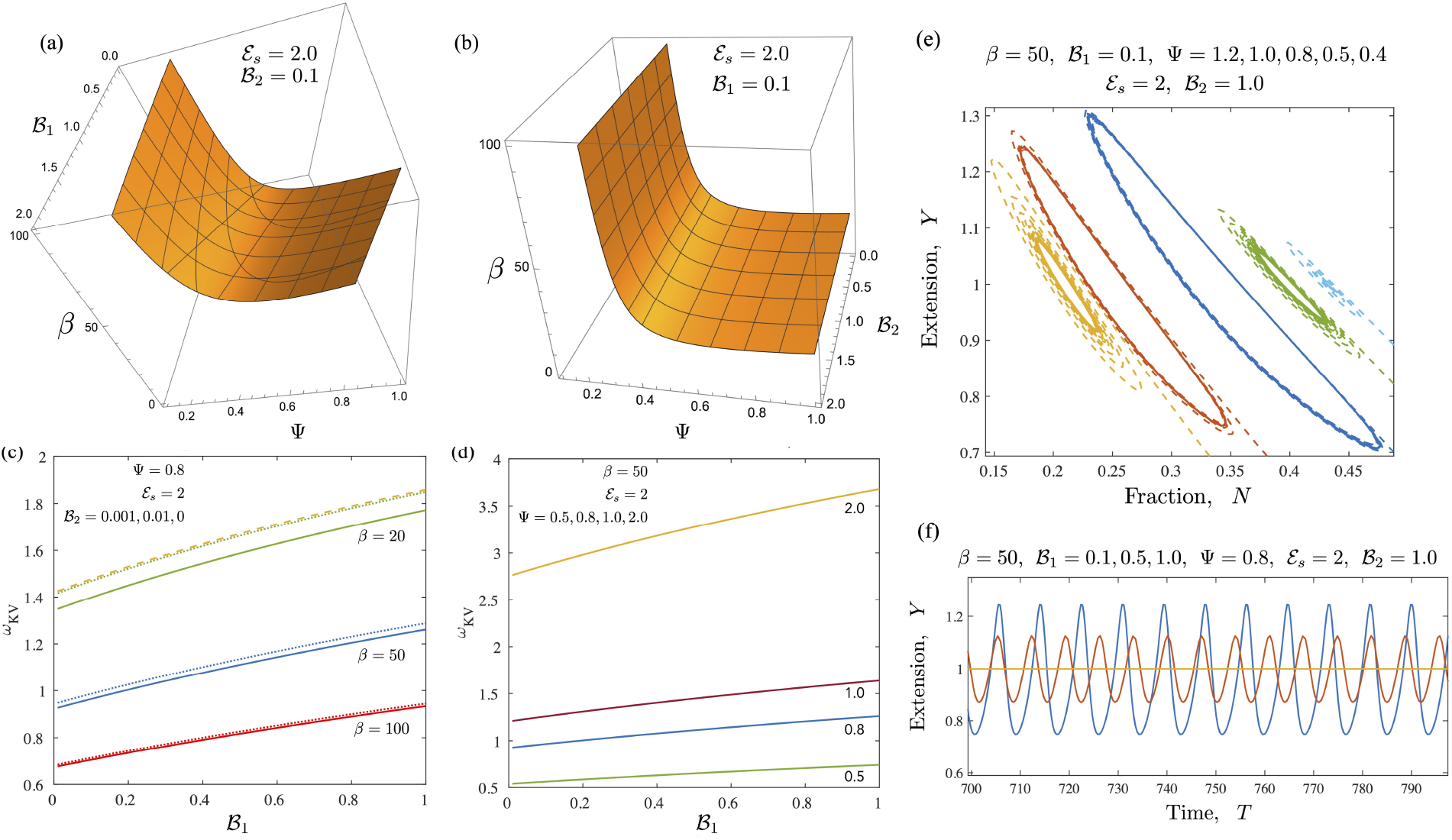
Results for the case of a Kelvin-Voigt medium. (c,d) Neutral stability curves from the linear stability analysis shown here for the Newtonian case when ℱ= cosh (*ε*_*s*_*Y*). Typically there are critical values of Ψ (for fixed *β*), or of *β* (for fixed Ψ) below which the base state is always stable. As plotted, the region on the concave side of the surface is linearly unstable. (c) & (d) Plots of the emergent frequencies (at onset) corresponding to equation (12), also obtained from the linear stability results. (e) & (f) Time-dependent long-time solutions obtained from a full non-linear solution to the ode system. In (e) we show plase-plots in *Y* − *N* space that demonstrates how see oscillations exist only for a range of Ψ values (with other parameters fixed). Subplot (f) demonstrates how increasing ℬ_1_ increases the frequency, and reduces the amplitude of the oscillations. Eventually at large values there is no instability.

**FIG. 6.**
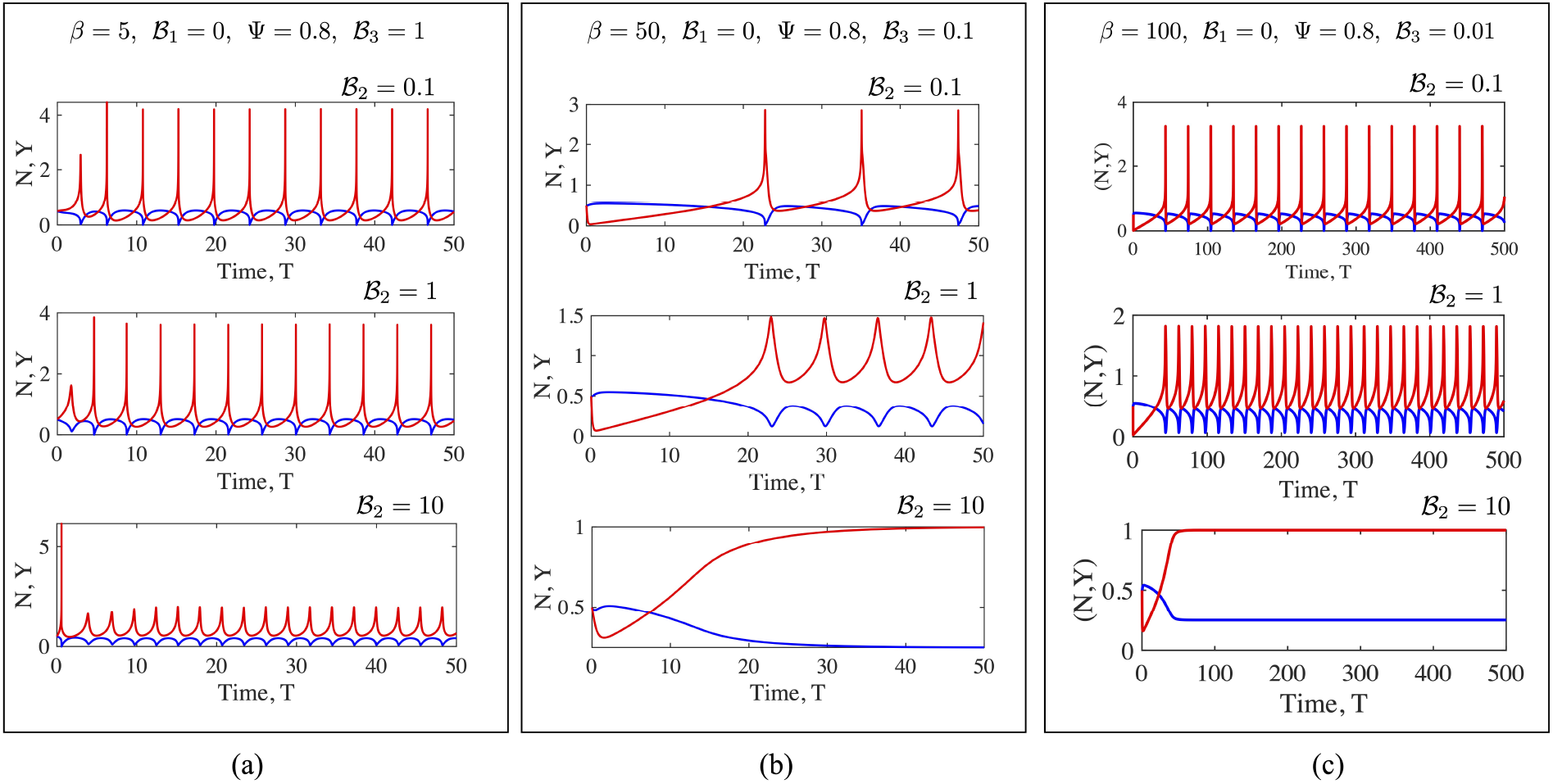
Fully non-linear solutions for the Maxwell medium. Here to focus on the rheological parameters ℬ_2_ and ℬ_3_, we study solutions for constant Ψ (= 0.8) with ℱ = cosh (*ε*_*s*_*Y*) (here *ε*_*s*_ = 2.0 for all cases). Results are shown for increasing values of *β* (from left to right) and increasing values of ℬ_2_ (top to bottom).

Let us start with the two reference cases first. For the case of no fluid drag, as anticipated by the linear stability analysis, oscillations can still exist. In this case the active energy input into the system by continuous motor activity is eventually dissipated away by internal motor friction with the anchoring spring as well as the passive links serving as intermediate storage agents. The internal friction while not explicitly stated is inherent in the assumption that motors that just detach eventually go back to their rest length. This loss of energy is associated with internal frictional mechanisms within the motor aggregate. We note that oscillations can feature very sharp gradients in *Y*. Over a cycle, the attached motor fraction *N* can become very small reaching relative values of *O*(10^−4^) as seen in Figure 4(a) (albeit for non-zero ℬ_2_ = 10^−3^). Also rather large values of *Y* are attained relative to the stationary state (*Y*_0_ = 1). Increasing *ε*_*s*_ enhances these features. When external viscosity starts to play a role and becomes comparable to motor friction, the parameter range over which instability is exhibited becomes smaller. This is seen in Figure 4(c) where increasing ℬ_2_ is seen to push the neutral stability envelope to the right. When at a point where oscillations are the stable state, increasing ℬ_2_ is seen to reduce variations in (*N, Y*) (evident when plotted as a phase-plot, Figure 4(a)). At For very large viscosity, oscillations become damped. Examination of the time-dependent behavior of *N* and *Y* just before and just after criticality confirms that the bifurcation is supercritical [37], with oscillations beginning with very small amplitudes. The farther the parameters are from their critical values, the larger the amplitude as is expected for a Hopf-Androvov-Poincare bifurcation.

Introducing viscoelastic effects changes the dynamics in a variety of ways. When the ambient medium follows Kelvin-Voigt like response (a reasonable simple approximation to a gel), the ability to store energy increases for ℬ_1_ *>* 0. When 0 < ℬ_2_ ≪ 1, and with increasing ℬ_1_ surprisingly increases the minimum value of Ψ (for fixed *β*) or the minimum value of *β* (for fixed Ψ) for the onset of oscillations. There is also a significant effect on the frequency on emergent solutions that we have analytically derived. Let us consider what happens when the motor parameters Ψ, *ε*_*s*_ and *β* are held fixed, while ℬ_1_ and _2_ are allowed to vary (Figure 5(c)). We note modest increases in the frequency with *lower* values of *β* yielding larger increments with increasing ℬ_1_ with ℬ_1_ = 0 reducing to the case of a pure Newtonian fluid medium. More dramatic increases in the frequency are seen when the kinetic parameter Ψ is allowed to vary and attain values larger than 1. Figure 5(d) illustrates a case with fixed ℬ_2_ where changing Ψ from 0.5 to 2.0 and simultaneously changing ℬ_1_ from 0 to 1 results in the frequency increasing by nearly a factor of 7. The amplitude of the oscillations however decreases with increasing ℬ_1_ with oscillations vanishing at sufficiently large values. Taken together, these results suggest that tuning the rheology of the fluid by adjusting ℬ_1_ allows for modest increases in frequencies with reduced solution amplitudes – both of these are consistent with experimental observations in more complicated ciliary and flagellar systems [9].

Numerical solutions for oscillations in a Maxwell fluid indicate that the onset of oscillations and frequencies may be more readily tuned than for the Kelvin-Voigt case. Oscillations are enabled at even *O*(1) values of *β* provided ℬ_3_ = *O*(1); with these oscillations persisting for even large values of the viscosity ℬ_2_. For fixed motor kinetics, in general, the higher the value of *β*, the lower ℬ_3_ needs to be to sustain oscillations. With Ψ, *β* and ℬ_1_ fixed, oscillations decrease in amplitude but increase in. frequency as ℬ_2_ increases. We hypothesize that the tuning of oscillations is made possible by changing the value of ℬ_3_ which is the ratio of the relaxation time for the Maxwell fluid and the mean time a motor is attached to the filament and is able to animate the filament. The viscosity on the other hand controls the energy dissipate rate and limits the amplitude of the oscillations. Figure 6 that shows sample limit cycles seen for various parameter values. For ℬ_3_ = 1 and keeping parameters Ψ = 0.8, *ε*_*s*_ = 2, ℬ_1_ = 0 fixed and restricting 0 < *ℬ*_2_ < 1, oscillations are seen for *β* = 5 but not for *β* = 50. Decreasing ℬ_3_ to 0.1 however moves the steady state to the linear stable phase space and thus oscillations are seen.

We note here that the elastic and viscous responses of for a viscoelastic fluid made by adding polymers to a Newtonian solvent fluid may depend on concentration differently. Thus in a polymeric fluid the ratio ℬ_1_/ℬ_2_ may scale with concentration with different exponents. Furthermore, the rheological behavior of a real polymeric fluid is more complex than that we have modeled. Nonetheless, the results here provide a foundation for more comprehensive analysis using more realistic models such as a retarded expansion for the stress-strain relationship or an anti-Zener like polymeric model (see Figure 1(c)).

## V. SUMMARY AND CONCLUSIONS

Here, we studied a minimal system comprised of an active bio-filament segment working against an elastic spring and immersed in model complex materials. A mean-field macroscale continuum description of filament-motor interactions was employed based on a cross-bridge model – however, other models can be readily used instead (see [35] for a general model that allows motors to undergo initial conformations as well as move relative to the filament). We used a combination of analytical and numerical methods to extract the conditions under which rheology of the fluid may be exploited to both shift the onset of instabilities, and modulate ensuing frequencies and amplitudes. We found that steady base states yield to stable oscillatory states or limit cycles. These solutions arise via supercritical Hopf-Andronov-Poincare’ bifurcations with exponentially damped oscillation change to small limit cycle oscillation about the steady state. Linear stability results were validated and complemented with full time-dependent solutions that provide an indication of the manner in which the amplitude of these oscillations changes.

The governing equations we utilized employed simplifications that allow for analytical and independent exploration of fluid viscoelastic properties (both its effective viscosity and elasticity, measured appropriately and encapsulated in parameters ℬ_1_ to ℬ_4_), motor kinetics and specifically the ratio of detachment and attachment constants (via parameter Ψ), effective importance of elastic (restoring) and viscous (dissipative) forces that includes possible coupling between the filament and its neighbors or anchoring substrates (through ratios ℬ_1_ and ℬ_2_), the ratio of viscoelastic relaxation times and the motor kinetic time-scales, and the strain dependent motor detachment rate (parameter 𝒮_*s*_). A further advantage of the simple model we have proposed is highlighted by examining the steady state solutions given by (5). We note that the steady state motor extension *Y*_0_ and the steady state fraction of attached motors *N*_0_ are *independent* of the rheological characteristics of the bulk medium. Thus coupling between the fluid and the motor aggregate is purely dynamical in nature. In other words, the aggregate is pre-strained internally by motor activity; but this pre-strained state is independent of the ambient fluid. It is thus possible to identify clearly changes in the dynamical state of the aggregate induced purely due to viscoelastic contributions, and to compare these with the results from the Newtonian case.

A shortcoming in our minimalist approach is that in the process of formulating analytically tractable equations, complex biochemical and biomechanical aspects are highly simplified. We have not considered noise and stochastic aspects of the problem that may be important for small to moderate values of *β* and/or low attachment rates [35]. While our model model does not include motor driven filament bending which dominates in oscillatory dynamics of biological and synthetically devised ciliary structures, reduced dimensional and spatially dependent versions have been used to investigate oscillations of cilia in Newtonian fluids [8, 9, 27, 29, 36]. However, ciliary or flagellar motion in viscoelastic fluids remains an open challenge. In this context, the model here may provide a foundation for more detailed computational and theoretical studies.

Our results are also relevant to understanding the rheology and dynamical response of biological and synthetic muscles, and in devising actuators for soft-robots [38–41]. In general, active responses to imposed external stimuli or perturbations are analyzed using an effective frequency-dependent dynamic moduli in these systems. The minimal approach we have used here - modeling the ambient fluid in terms of minimal models with spring-like and dashpot-like elements – provide a clear means to evaluate these dynamic response functions (or transfer functions).

## References

[1] J. Howard (2001) Mechanics of motor proteins and the cytoskeleton, Sinauer associates, Sunderland.

[2] S. W. Grill, K. Kruse and F. Jülicher (2005) Theory of mitotic spindle oscillations. Phys. Rev. Lett., 94(10): 108104.

[3] E. Bizzi, W. Chapple and N. Hogan (1982) Mechanical properties of muscles: implications for motor control. Trends Neurosci. 5, 395–398. doi: 10.1016/0166-2236(82)90221-1.

[4] N. Hogan (1984) Adaptive control of mechanical impedance by coactivation of antagonist muscles. IEEE Trans. Automatic Control 29, 681–690. doi: 10.1109/TAC.1984.1103644.

[5] S. Walcott (2014) Muscle activation described with a differential equation model for large ensembles of locally coupled molecular motors. Phys. Rev. E 90:042717. doi: 10.1103/PhysRevE.90.042717.

[6] G. B. Witman, Introduction to cilia and flagella in ciliary and flagellar membranes (ed. R. A. Bloodgood), Plenum, New York, pp 1–30 (1990).

[7] K.E. Machin (1963) The control and synchronization of flagellar movement. Proc. Roy. Soc. B., 158(970), 88–104.

[8] C. J. Brokaw (1975) Molecular mechanism for oscillation in flagella and muscle. Proc. Natl. Acad. Sci. USA, 72(8): 3102–3106.

[9] B. Qin, A. Gopinath, J. Yang, J. J. Gollub and P. E. Arratia (2015) Flagellar Kinematics and Swimming of Algal Cells in Viscoelastic Fluids. Scientific Reports, 5, 9190.

[10] P. R. Sears, K. Thompson, M. R. Knowles and C. W. Davis (2013) Human airway ciliary dynamics. Am J Physiol Lung Cell Mol Physiol., 304:L170–L183.

[11] C. Kempeneers, C. Seaton and M. A. Chilvers (2017) Variation of ciliary beat pattern in three different beating planes in healthy subjects. Chest, 151: 993–1001.

[12] M. R. Knowles, M. W. Leigh, J. L. Carson, S. D. Davis, S. D. Dell, T. W. Ferkol et al. (2012) Genetic Disorders of Mucociliary Clearance Consortium. Mutations of DNAH11 in patients with primary ciliary dyskinesia with normal ciliary ultrastructure. Thorax, 67:433–441.

[13] J. F. Papon, L. Bassinet, G. Cariou-Patron, F. Zerah-Lancner, A. M. Vojtek, S. Blanchon S, et al. (2012) Quantitative analysis of ciliary beating in primary ciliary dyskinesia: a pilot study. Orphanet J Rare Dis, 7:78.

[14] B. Thomas, A. Rutman and C. O’Callaghan (2009) Disrupted ciliated epithelium shows slower ciliary beat frequency and increased dyskinesia. Eur Respir J., 34:401–404.

[15] C. Ringers, E. W. Olstad and N. Jurisch-Yaksi (2019) The role of motile cilia in the development and physiology of the nervous system. Phil. Trans. Roy. Soc. B., 375:20190156.

[16] T. Sanchez, D. Welch, D. Nicastro, Z. Dogic (2001) Cilia-like beating of active microtubule bundles. Science, 333(6041): 456–459.

[17] T. Sanchez, D. T. N. Chen, S. J. DeCamp, M. Heymann, Z. Dogic (2012) Spontaneous motion in hierarchically assembled active matter. Nature, 491(7424): 431–434.

[18] A. Vilfan and E. Frey (2005) Oscillations in molecular motor assemblies. J. Phys. Condens. Matter, 17(47):S3901-S3911.

[19] E. Evans and K. Ritchie (1997) Dynamic strength of molecular adhesion bonds. Biophys. J., 72(4):1541-1555.

[20] K. D. Nguyen, N. Sharma and M. Venkadesan (2018) Active Viscoelasticity of Sarcomeres. Front. Robot. AI 5:69. doi: 10.3389/frobt.2018.00069.

[21] A. Vilfan and T. Duke (2003) Synchronization of active mechanical oscillators by an inertial load. Phys. Rev. Lett., 91:114101.

[22] S. A. Endow and H. Higuchi (2000) A mutant of the motor protein kinesin that moves in both directions on microtubules. Nature, 406:913–916.

[23] M. Badoual, F. Julicher and J. Prost (2002) Bidirectional cooperative motion of molecular motors. Proc. Natl. Acad. Sci. USA, 99:6696–6701.

[24] S. W. Grill, P. Gonczy, E. H. Stelzer and A. A. Hyman (2001) Polarity controls forces governing asymmetric spindle positioning in the Caenorhabditis elegans embryo. Nature, 409:630–633.

[25] K. Colombo, S. W. Grill, R. J. Kimple, F. S. Willard, D. P. Siderovski and P. Gonczy P (2003) Translation of polarity cues into asymmetric spindle positioning in Caenorhabditis elegans embryos. Science,300:1957–1961.

[26] S. W. Grill, J. Howard, E. Schaffer, E. H. Stelzer and A.A. Hyman AA (2003) The distribution of active force generators controls mitotic spindle position. Science, 301:518–521.

[27] S. Camalet and F. Jülicher (2000) Generic aspects of axonemal beating. New J. Phys., 2: 24.1–24.23.

[28] C. Li, B. Qin, A. Gopinath, P. E. Arratia, B. Thomases and R. D. Guy (2017) Flagellar swimming in viscoelastic fluids: role of fluid stress revealed by simulations based on experimental data. Proc. Roy. Soc. Interface. 14(135), 20170289.

[29] E. Hamilton, N. Pelliciotta, L. Feriani and P. Cicuta (2019) Motile cilia hydrodynamics: entrainment versus synchronization when coupling through flow. Phil. Trans. Roy. Soc. B., 375:20190152.

[30] A. Sangani and A. Gopinath (2020) Elastohydrodynamical instabilities of active filaments, arrays and carpets analyzed using slender body theory bioRxiv 986596 (bioRxiv 2020.03.10.986596).

[31] S. Fatehiboroujeni, A. Gopinath and S. Goyal (2018) Nonlinear Oscillations Induced by Follower Forces in Prestressed Clamped Rods Subjected to Drag. J. Comp. Non. Dyn. 13 (12):121005.

[32] S. Fatehiboroujeni, A. Gopinath and S. Goyal (2018) Follower Forces in Pre-Stressed Fixed-Fixed Rods to Mimic Oscillatory Beating of Active Filaments. ASME DETC2018-85449, V006T09A033.

[33] R. Chelakkot, A. Gopinath, L. Mahadevan and M. F. Hagan (2014) Dynamics of connected self-propelled Brownian particles. Proc. Roy. Soc. Interface, 11(92), 20130884.

[34] Y. Fily, P. Subramaniam, T. M. Scheider, R. Chelakkot and A. Gopinath (2020) Buckling instabilities and spatiotemporal dynamics of active elastic filaments. J. Roy. Soc. Interface 17 (165):20190764.

[35] A. Gopinath, R. Chelakkot and L. Mahadevan (2020) Filament extensibility and shear stiffening control persistence of strain and loss of coherence in cross-linked motor-filament assemblies. https://doi.org/10.1101/423582

[36] B. Chakrabarti and D. Saintillan (2019) Spontaneous oscillations, beating patterns, and hydrodynamics of active microfilaments. Phys. Rev. Fluids 4, 043102.

[37] S. H. Strogatz (1994) Nonlinear Dynamics and Chaos: With Applications to Physics, Biology, Chemistry, and Engineering. Reading, Mass: Addison-Wesley.

[38] I. A. Anderson, T. A. Gisby, T. G. McKay, B.M. O’Brien and E. P. Calius (2012) Multi-functional dielectric elastomer artificial muscles for soft and smart machines. J. Appl. Phys. 112:041101. doi: 10.1063/1.4740023.

[39] S. P. Buerger and N. Hogan (2007) Complementary stability and loop shaping for improved human–robot interaction. IEEE Transactions on Robotics 23, 232–244. doi: 10.1109/TRO.2007.892229.

[40] N. T. George, T. C. Irving, C. D. Williams and T. L. Daniel (2013) The cross-bridge spring: can cool muscles store elastic energy? Science 340, 1217–1220. doi: 10.1126/science.1229573.

[41] L. Hines, K. Petersen, G. Z. Lum and M. Sitti (2017) Soft actuators for small-scale robotics. Adv. Mat. 29, 1–43. doi: 10.1002/adma.201603483

